# Attenuation of the CpG Island Methylator Phenotype and lack of WNT signaling activation restrains *Kras* mutant intestinal neoplasia

**DOI:** 10.1101/2023.12.21.572936

**Authors:** Lochlan Fennell, Simon Tria, Cheng Liu, Alexandra Kane, Diane McKeone, Jennifer Borowsky, Lu Chai, Sarron Randall-Demllo, Catherine Bond, Barbara Leggett, Vicki Whitehall

**Affiliations:** Oncogenic Aging Laboratory; Biomedicine Discovery Institute, Monash University, Clayton, Australia; Conjoint Gastroenterology Laboratory; Cell and Molecular Biology Department, QIMR Berghofer Medical Research Institute, Herston, Australia; Faculty of Medicine, University of Queensland, St Lucia, Australia; Conjoint Internal Medical Laboratories; Pathology Queensland, Brisbane, Australia; Envoi Specialist Pathologists; Kelvin Grove, Australia; Department of Gastroenterology and Hepatology; Royal Brisbane and Women’s Hospital, Herston, Australia

**Author notes:** Address for Correspondence: Lochlan Fennell Level 3, STRIP2 - Monash University 19 Innovation Walk, Clayton, 3168 Australia. These authors contributed equally to this work (Lochlan Fennell, Simon Tria).

## Abstract

**Background:** Serrated neoplasia accounts for ∼25% of colorectal cancer. These cancers arise from serrated precursor lesions. Hyperplastic polyps initiated by either *BRAF* or *KRAS* mutation activating MAPK signalling are common, but premalignant sessile serrated lesions with *KRAS* mutation are rare. Here, we model *Kras* and *Braf* mutant neoplasia *in vivo* to compare histological, gene expression and DNA methylation manifestations associated with activation of these oncogenes.

**Methods:** We employ cre-recombinase dependent *Braf*^V637^ and *Kras^G12D^* murine models, and cross animals with those bearing the Villin-Cre^ERT2^ transgene to direct temporospatial activation of these oncogenes to the murine intestine. We examine histology, and genome-scale DNA methylation and gene expression via reduced representation bisulphite sequencing and RNA-Seq, respectively. We performed differential gene expression, methylation and pathways analysis to identify oncogene specific alterations.

**Results:** Prolonged exposure to oncogenic *Braf* is associated with a time-dependent accumulation of murine serrated precursors (P=3x10^-10^) and advanced murine serrated lesions and invasive cancer (8x10^-8^). *Kras* mutant animals acquire fewer precursor lesions (P=0.06) and have a significantly lower probability of developing advanced serrated lesions (P=0.004). *Braf* and *Kras* mutant animals develop pronounced hyperplasia, however the severity is significantly less in *Kras* mutant animals. *Kras* mutant advanced serrated lesions rarely develop aberrant WNT signaling activation (1/23). Gene expression profiling showed divergent transcriptomic profiles between *Braf* and *Kras* mutant intestines, with the former overexpressing genes associated with immune and inflammatory signaling. Deconvolution analysis revealed a comparably higher macrophage infiltrate (P=0.025) and upregulation of M1 macrophage gene sets in the *Braf* mutant intestine (P=0.0008), contributing to chronic inflammatory signalling. Both *Kras* and *Braf* mutations lead to accumulation of substantial temporal DNA methylation alterations, however a subset of CpG sites (1,306) show an attenuated rate of DNA methylation accumulation in the *Kras* mutant intestine compared with *Braf* mutant animals.

**Conclusions:** In this study, we show that *Kras* mutation can induce serrated intestinal neoplasia, however the latency period and penetrance is significantly lower when compared with *Braf* mutation. Aberrant WNT signalling is common in lesions arising in the context of *Braf* mutation, but rare in *Kras* mutant neoplasms. We show marked transcriptomic disparities between these models, with a tendency for the *Braf* mutant intestine to upregulate immunological processes. Our DNA methylation analysis reveals an attenuated CIMP-like phenotype that is specific to the *Kras* mutant intestine, consistent with our previous works in humans. These data have significant implications for our understanding of how MAPK-induced neoplasia develops within the intestine.

**Synopsis:** *BRAF* and *KRAS* mutant hyperplastic polyps have disparate malignant potential and the reason for this is unclear given both oncogenes activate MAPK signalling. We show that the DNA methylation alterations that follow *Kras* mutation are attenuated and that hyperactivation of WNT signaling is rare, providing a molecular mechanism that restrains malignant transformation.

## Introduction

Serrated colorectal neoplasia is now accepted to be the molecular pathway underpinning ∼25% of colorectal cancers. These cancers are associated with female gender, proximal colonic location and older age. Serrated colorectal cancers are also molecularly distinct from conventional pathway cancers. They are driven by *BRAF* mutation, rarely display chromosomal instability, and ∼70% have a hyper-mutator phenotype. Serrated pathway colorectal cancers arise from sessile serrated lesions (SSLs) and less commonly traditional serrated adenomas (TSAs). SSLs are strongly associated with *BRAF* mutation, and rarely possess *KRAS* mutation. TSAs, by contrast, can be initiated by *BRAF* or *KRAS* mutation (1,2) but are very uncommon. Both *BRAF* and *KRAS* reside in the MAPKinase signalling pathway, and their mutation results in increased proliferation, diminished apoptosis, and increased cell growth (3). It is not clear why *KRAS* mutation does not appear to be sufficient to initiate serrated neoplasia via SSLs.

The CpG Island Methylator Phenotype (CIMP) is an important component of *BRAF* mutant serrated neoplasia. Several studies have implicated *BRAF* mutation directly in causing this phenotype. It is possible that *KRAS* mutation does not induce the degree of DNA hypermethylation required for progression of sessile serrated lesions. This is supported by DNA methylation analysis of hyperplastic polyps. *BRAF* mutant hyperplastic polyps tend to have a microvesicular histological appearance (4), and can harbour molecular features that are similar to sessile serrated lesions, such CIMP (5). In contrast, most *KRAS* mutant hyperplastic polyps have a goblet-cell rich histological appearance, and rarely display CIMP (5). These findings might indicate that the lack of CIMP is a rate limiting step in serrated neoplasia via sessile serrated lesions. Another critical step in the transition from SSL to cancer is the acquisition of molecular changes which increase WNT signalling (6).

Since both *KRAS* and *BRAF* act within the MAPKinase pathway such that mutations are mutually exclusive, it is not clear why serrated polyps initiated by *KRAS* mutation do not appear to have the same malignant potential as their *BRAF* mutant counterparts. Using a *Kras^LSL-G12D^* mouse model, Feng et al (7) reported intestinal hyperplasia and an alteration in the cellular composition of intestinal crypts, but did not observe pre-malignant lesion formation. They attributed this to a lack in the expansion of the presumptive stem pool. This, and previous studies using *Kras* mutant mice, were medium term studies. It is possible that lesion formation is protracted in these models, and this has reduced the likelihood of detecting them.

In this study, we have performed a comprehensive, long-term comparative analysis of the histological, gene expression and DNA methylation alterations induced by oncogenic *Braf* and *Kras in vivo* to shed light on the differential role of MAPK mutations in the colorectum. We report substantial transcriptomic, epigenetic and histological differences underpinning *Braf* and *Kras* induced neoplasia, highlighting the divergent nature of the molecular evolution of neoplasia in these contexts.

## Methods

### Murine models of serrated neoplasia

To contrast the effects of *Braf* and *Kras* mutation on the murine intestine we employed two transgenic murine models. The *Braf^V637^* and *Kras^G12D-LSL^* models are cre-inducible and facilitate the expression of the oncogenic *Braf^V637E^* and *Kras^G12D^* alleles, respectively. These are the murine analogues of the frequently mutated human *BRAF^V600E^* and *KRAS^G12D^* alleles, and recapitulate the constitutive oncogenic signaling that follows these mutations. These animals were crossed with Villin-Cre^ERT2^ mice, providing efficient spatiotemporal control over the recombination of the oncogenic alleles via a single intraperitoneal injection of tamoxifen (75mg/kg).

### Immunohistochemistry

Immunohistochemistry was performed to assess the protein expression of β-Catenin and Ki67. For Ki67, 4µm tissue sections were affixed to charged slides, dewaxed, and rehydrated. Peroxidase activity was blocked by incubating sections in 2% hydrogen peroxide in tris-buffered saline for 10 minutes. Antigens were retrieved by incubating sections in pH 6.0 Epitope Retrieval Solution (Dako, USA) at 125⁰C for 3 minutes in a decloaking chamber. Sections were incubated in Background Sniper + 1% BSA (Biocare Medical, USA) to prevent spurious staining. Antibody was applied to sections for 60 minutes (1:100 dilution in Da Vinci Green (Biocare Medical); Ab Cat #ab16667). Stains were developed by incubating sections in MACH1 Rabbit HRP for 30 minutes and betazoid DAB for 5 minutes, and counterstained with Haematoxylin. Slides were scanned using an Aperio AT2 slide scanning instrument and analysed using QuPath using default cell segmentation and positive detection settings. β-Catenin immunohistochemistry was performed as per Kane et al 2020. Briefly, 4uM sections were mounted on charged slides, dewaxed with xylol and rehydrated via the descending graded alcohol method. Peroxidase activity was blocked as outlined above. To retrieve antigens, slides were incubated in Dako antigen retrieval solution (pH 6.0) for 5 minutes at 125⁰C in a decloaking chamber. Non-specific binding was blocked using Background Sniper + 2.0% BSA (Biocare Medical, USA). Sections were then incubated with an anti-human β-Catenin monoclonal antibody diluted 1:500 in 2% BSA in TBS (Ab cat 32572) overnight at room temperature. Stains were developed by incubating sections in MACH2 Rabbit HRP for 30 minutes and betazoid DAB for 5 minutes, and counterstained with Haematoxylin. Sections were graded by an expert gastrointestinal pathologist (JB) as per Kane et al (8), which included an assessment of staining intensity, extent and localization across lesions.

### Nucleic acid extractions and next generation sequencing library preparations

At necropsy, fresh tissue samples were harvested from the small intestine and snap frozen in liquid nitrogen. DNA and RNA were extracted from ∼30mg of fresh-frozen tissue using the AllPrep DNA/RNA/Protein mini kit as per the manufacturer’s instructions. Sample integrity and quantity was assessed using the TapeStation platform (Agilent Technologies, USA). DNA methylation was assessed via reduced representation bisulphite sequencing (RRBS). Libraries were generated using the Ovation RRBS Methyl-Seq library preparation kit (Tecan Genomics, USA) as previously reported (Fennell et al, 2021). Transcript expression was assessed via RNA-Seq using the Illumina Stranded Total RNA with RiboZero Plus kit (Illumina, USA) as per the manufacturers protocol.

### RNA-Sequencing data analysis

RNA-Sequencing data was processed using nfcore rnaseq (9) nextflow pipeline (v3.0). This pipeline performs adaptor and quality trimming with Trim Galore! (v0.6.6), aligns reads to GRCm38 using STAR (10) (v2.6.1d), and quantifies genes and transcripts using Salmon(11) (v1.4.0). The pipeline generates various quality control metrics using tools including RSeQC(12) (v3.0.1), Qualimap(13) (v2.2.2), FastQC (v0.11.9), dupRadar(14) (v1.18.0), and Preseq (v2.0.3). Differential gene expression analysis was performed using DeSeq2 (15) (v3.13). To estimate and compare cell type proportions we deconvoluted the bulk RNAseq data using ImmuneCC, a reference-based deconvolution method, and the standard immune signature matrix. To prepare the data for this analysis we performed TMM normalization on raw counts using EdgeR (v3.4.2). Data was kept in linear space as per best practice for bulk deconvolution.

### Reduced Representation Bisulphite Sequencing data analysis

Reduced representation bisulphite sequencing reads were trimmed to remove adaptors and poor quality sequences using Trim Galore! (v0.6.6). NuGen diversity adaptors were removed using the custom ‘trimRRBSdiversityAdaptCusomters.py’ script(16). Reads were then aligned and methylation calls generated using the methylseq nf-core pipeline (v2.3.0). This pipeline employs Bismark for methylation aware sequence alignment and extraction, and qualimap and preseq for generating quality control metrics. Principal components analysis was performed using the ‘prcomp’ R function. Differential DNA methylation analysis was performed using Limma (v3.56.2) and differential methylation results were considered significant if the absolute methylation difference in contrasts was > 20 % and the FDR < 0.05. The methylGSA (v1.18.0) method was employed for gene ontology enrichment analysis using the robust rank aggregation method with CpGs mapping to promoters (distance to transcription start site: -1500 to +100bp) as input. This method adjusts for the extreme biases that exist when performing gene set analysis on DNA methylation data by virtue of the unequal distribution of CpGs in the promoters of protein coding genes. To assess age associated DNA methylation we performed linear regression analysis using the ‘lm’ R function. We then assessed for mutation specific differences in age-associated DNA methylation by analysis of covariance.

### Statistical Analyses

All statistical analyses were performed using R (v4.0.3). Results were reported as significant if p < 0.05. P values were corrected using the false discovery rate method, where appropriate.

## Results

### Oncogenic Braf and Kras mutation induces differentially penetrant murine serrated neoplasia

*BRAF* and *KRAS* mutation are common mutational events in human serrated colorectal neoplasia. To test whether mutation of *BRAF* or *KRAS* is sufficient to initiate serrated neoplasia we employed the spatiotemporally inducible *Braf^V637^* and *Kras^G12D^* murine models (see methods for more detail).

In *Braf^V637^* mice, we observe a steady and significant temporal accumulation of murine serrated precursors (R^2^=0.56, P= 3x10^-10^; Figure 1A). These lesions display crypt dilatation and mucinous differentiation, consistent with previous studies (17). *Kras* mutant mice, by contrast, accumulated lesions at a much slower rate (R^2^ = 0.22, P=0.04; Figure 1A) and were rare even in those exposed to *Kras* mutation for 14-18 months.

**Figure 1:**
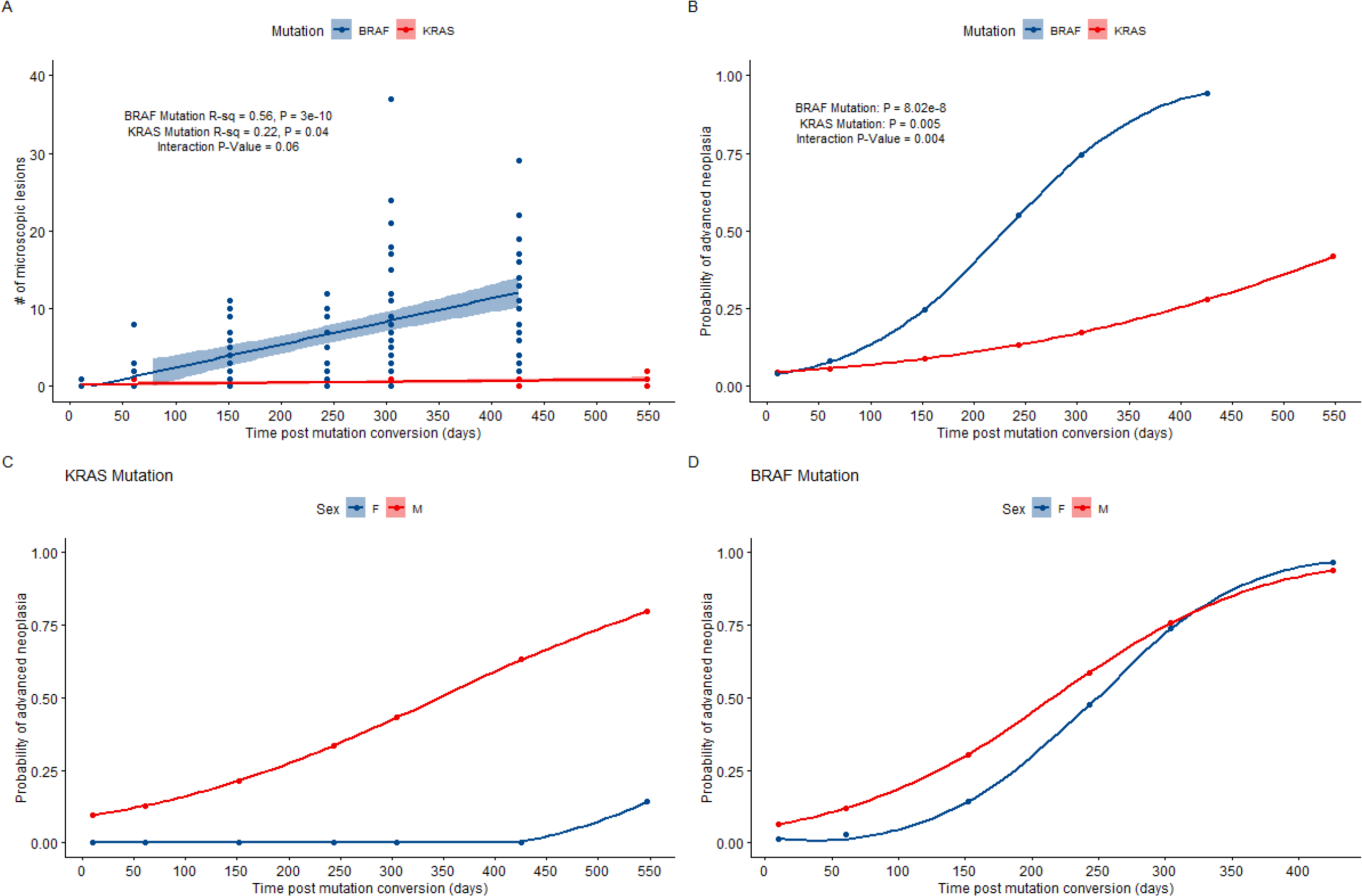
A) Number of microscopic lesions detected in *Braf* and *Kras* mutant animals aged from 10 days to 18 months. B) The probability of advanced serrated neoplasia developing as a function of age and time exposed to each oncogene from wean. C & D) Sex specific probability of advanced serrated neoplasia as a function age and time exposed to each oncogene from wean.

Both models developed advanced serrated lesions, which display overt cytological dysplasia; however the frequency and rate at which these lesions developed differed substantially between the models. The probability of *Braf* mutant animals developing an advanced serrated lesion increased substantially from ∼25% in five-month animals to >95% in 14 month animals. In contrast, the probability of *Kras* mutant animals developing advanced serrated neoplasia increased significantly more slowly with age (Logistic regression interaction model P=0.004; Figure 1B). At the 14 month time point, the probability of a *Kras* mutant animal developing advanced serrated neoplasia was 66% lower than age matched *Braf* mutant animals (28% vs 94%; Figure 1B). Therefore, both *Braf* and *Kras* mutation can induce serrated neoplasia, however the *Braf* mutation has a significantly higher penetrance.

*BRAF* mutant human cancers tend to occur more frequently in female individuals, and *KRAS* mutant cancers have a preponderance for males. Thus, we next sought to assess whether there were any gender biases present in our murine models. *Braf-*induced advanced serrated neoplasia at the same frequency in male and female animals (Figure 1D). However, *Kras-*induced serrated neoplasia occurred almost exclusively in male animals (P<0.0001), with only one *Kras* mutantfemaleanimal developing an advanced serrated lesion in the entire study. When segregated by gender, *Kras* mutant male animals had a 79% chance of developing advanced serrated neoplasia at 18 months (Figure 1).

### *Kras* mutation induces less severe hyperplasia, but greater crypt proliferation when compared with *Braf* mutation

To examine whether proliferative activity might underpin the histological differences we observe in the manifestations of both mutations, we evaluated the extent of hyperplasia in the *Braf* and *Kras* mutant intestines. First, we examined the length of the small intestine of all animals in the study. These measurements were recorded at necropsy. At all timepoints assessed in both models (10 days to 14 months), the *Braf* mutated intestine was significantly longer as compared with *Kras* mutant animals (ANOVA P = 0.0002). Male *Kras* mutant animals had significantly longer small intestines than female *Kras* mutant animals (ANOVA P = 6.83x10^6^, mean difference 3.63cm), however there was no specific interaction between sex and intestine length in *Braf* mutant animals (ANOVA P = 0.18). Next, we measured the length of the small intestinal villi of a subset of animals (15 villus per animal). In keeping, we observed that the length of the villi of *Kras* mutant animals was significantly shorter when compared with age-matched *Braf* mutant mice (Figure 2A). We did not observe any sex specific differences in villus length in either setting. These data indicate that *Braf* mutation induces greater hyperplasia when compared with *Kras* mutation.

**Figure 2:**
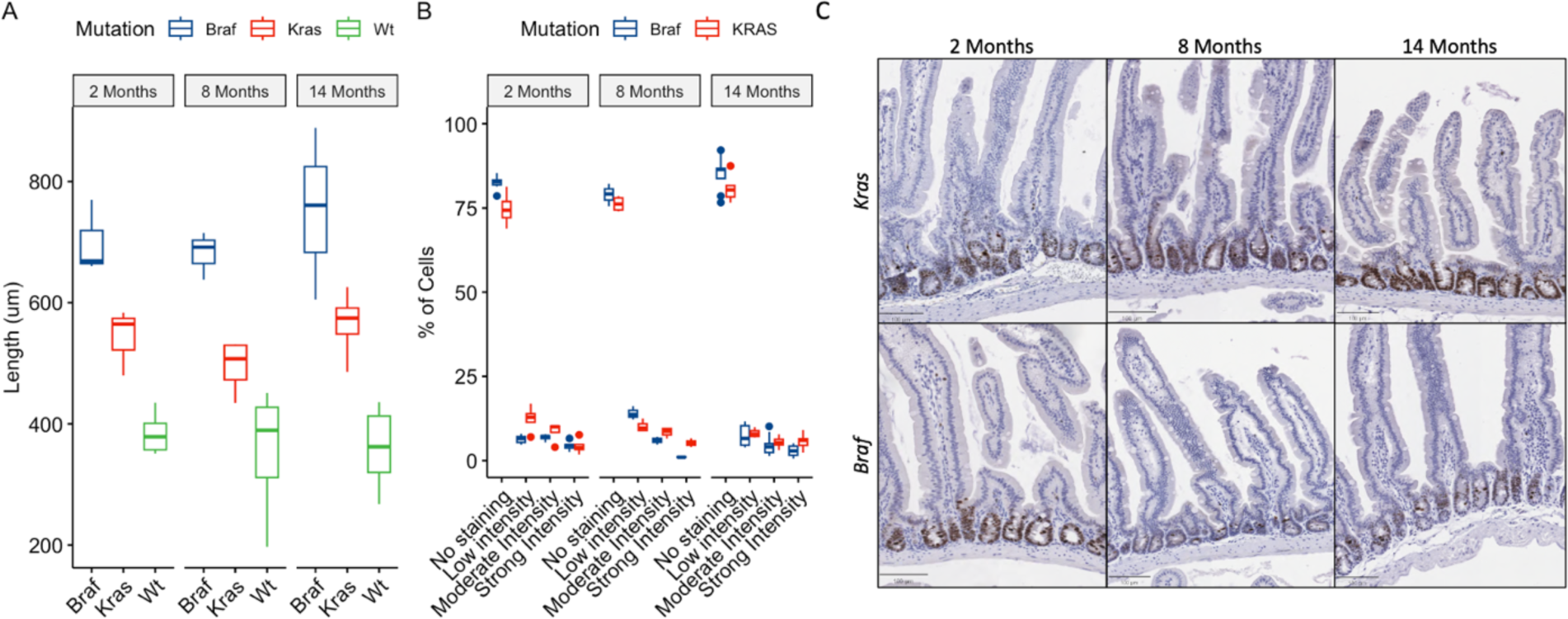
A) Mean Villus length of *Braf, Kras* and WT animals at two, eight and fourteen months. For each animal 15 villi were measured by a blinded operator. B) Proliferative activity in the *Braf and Kras* intestines as measured by Ki67 immunostaining. (*Braf:* n= 7, 3 and 12 for 2 months, eight months and fourteen months, respectively; *Kras:* n= 4, 4, and 6 for two, eight and fourteen months, respectively) C) Representative images of Ki67 staining in *Braf* and *Kras* mutant animals at two, eight and fourteen months.

Next, we sought to examine whether the more severe hyperplasia observed in *Braf* mutant animals might be explained by greater proliferative activity in the intestinal crypts. Ki67 accumulates in the nucleus during the cell cycle and is a surrogate marker of proliferation. We performed immunohistochemical analysis of intestinal sections to compare the proliferation of *Braf* and *Kras* mutant mice. At all time points, we observed significantly greater proliferation in the crypts of *Kras* mutant compared with *Braf* mutant animals (Figure 2B-C; Two-way ANOVA P=4.75x10^6^, 0.0013, and 0.024 at two, eight and fourteen months, respectively).

### WNT signaling hyperactivation is not a feature of Kras mutant serrated lesions

WNT signaling activation is a hallmark of colorectal neoplasia. We have previously shown that *Braf* mutant lesions acquire dysregulated WNT signalling via mutation of *CTNNB1,* which results in a concomitant translocation of β-catenin to the nucleus and activation of WNT-target genes (8,17). To test whether WNT signalling is similarly deregulated in *Kras* mutant serrated neoplasia, we performed immunohistochemical analysis on murine serrated lesions (n=16 animals, n=23 total lesions). Abnormal β-catenin staining was observed in 1/23 lesions assessed (Figure 3A). This lesion displayed weak focal nuclear β-catenin affecting <10% of nuclei in the lesion (Figure 3B). This lesion arose in a 14 month old animal. The remaining lesions displayed a normal membranous staining pattern. By contrast, we previously reported that 40% and 100% of *Braf* mutant mSLs had abnormal β-catenin staining 10 and 14 months, respectively (8). These data indicate that abnormal WNT signalling activation is not a feature of murine serrated lesions that arise in the setting of *Kras* mutation.

**Figure 3:**
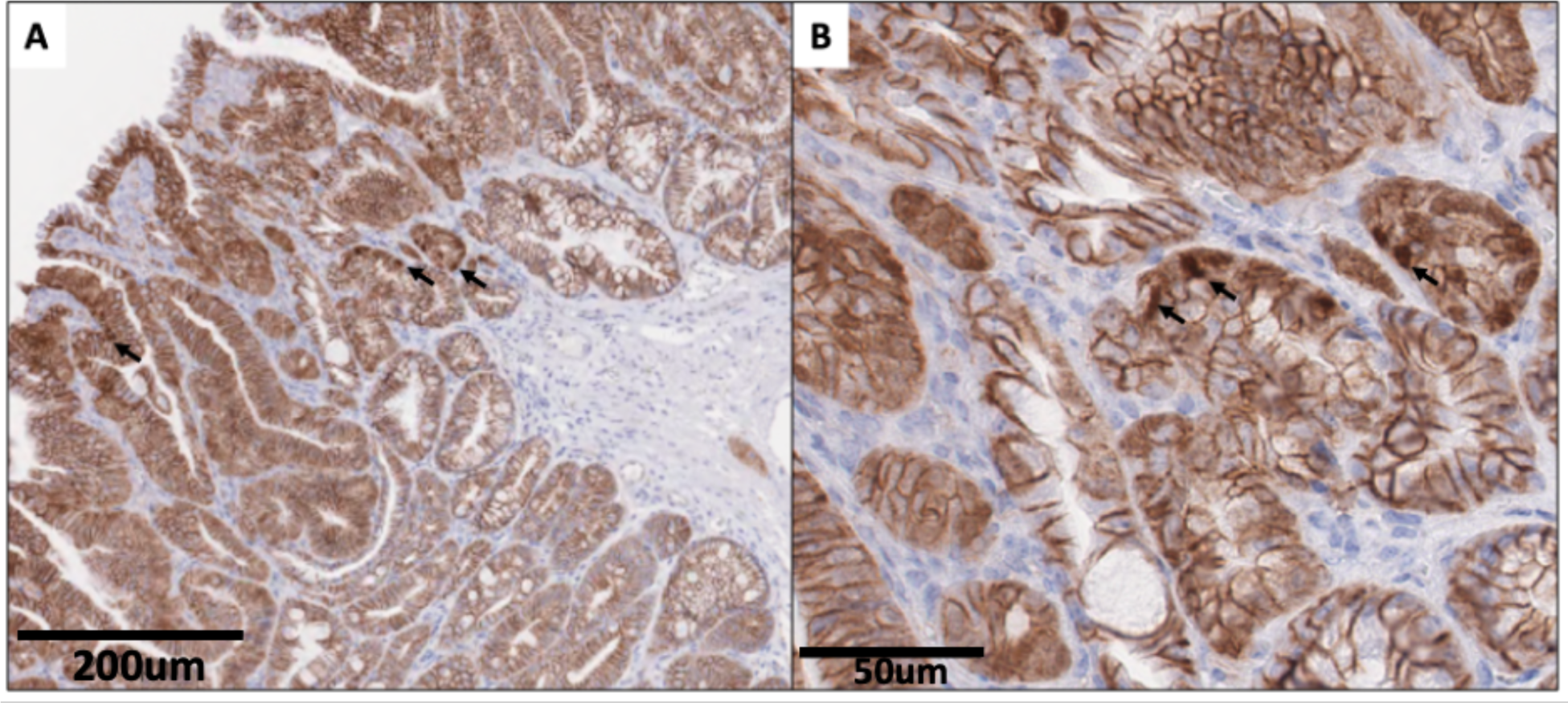
β-catenin immunohistochemistry showing focal nuclear localization of β-catenin (arrows) in the sole *Kras* mutant murine serrated lesion where abnormal β-catenin was identified. Typical membranous staining is observed throughout the remainder of the lesion. Brown staining: DAB Blue staining: Hematoxylin.

### Temporal divergence of the intestinal transcriptome in *Braf* and *Kras* mutant samples

Next we sought to examine whether different signaling pathways are activated upon mutation of *Braf* when compared with mutation of *Kras*. We performed RNAseq on animals *Kras* and *Braf* mutant intestinal samples at 2 months, 8 months, and 14 months post mutation activation (n=3 animals/genotype/timepoint).

Differential gene expression analysis revealed 148 differentially expressed genes between 2 month old *Kras* and *Braf* mutant samples (Absolute LogFC > 2, FDR < 0.05), of which 38 were overexpressed in *Kras* mutant samples, relative to *Braf* mutant samples, and 110 underexpressed (Table 1, Figure 4A). At the 8 month timepoint, we observe significantly altered gene expression in 1023 genes. In keeping, the majority (790 genes, Table 1) were underexpressed in *Kras* mutant samples, with 233 genes being expressed to a greater degree in *Kras* mutant samples (Table 1, Figure 4A). By 14 months, 557 genes were differentially regulated, with 328 genes being underexpressed and 229 overexpressed in *Kras* mutant samples relative to age-matched *Braf* mutant samples (Table 1, Figure 4)). These data indicate that some loci are immediately differentially regulated in response to either *Kras* or *Braf* mutation, while others require sustained oncogenic exposure to manifest.

**Figure 4:**
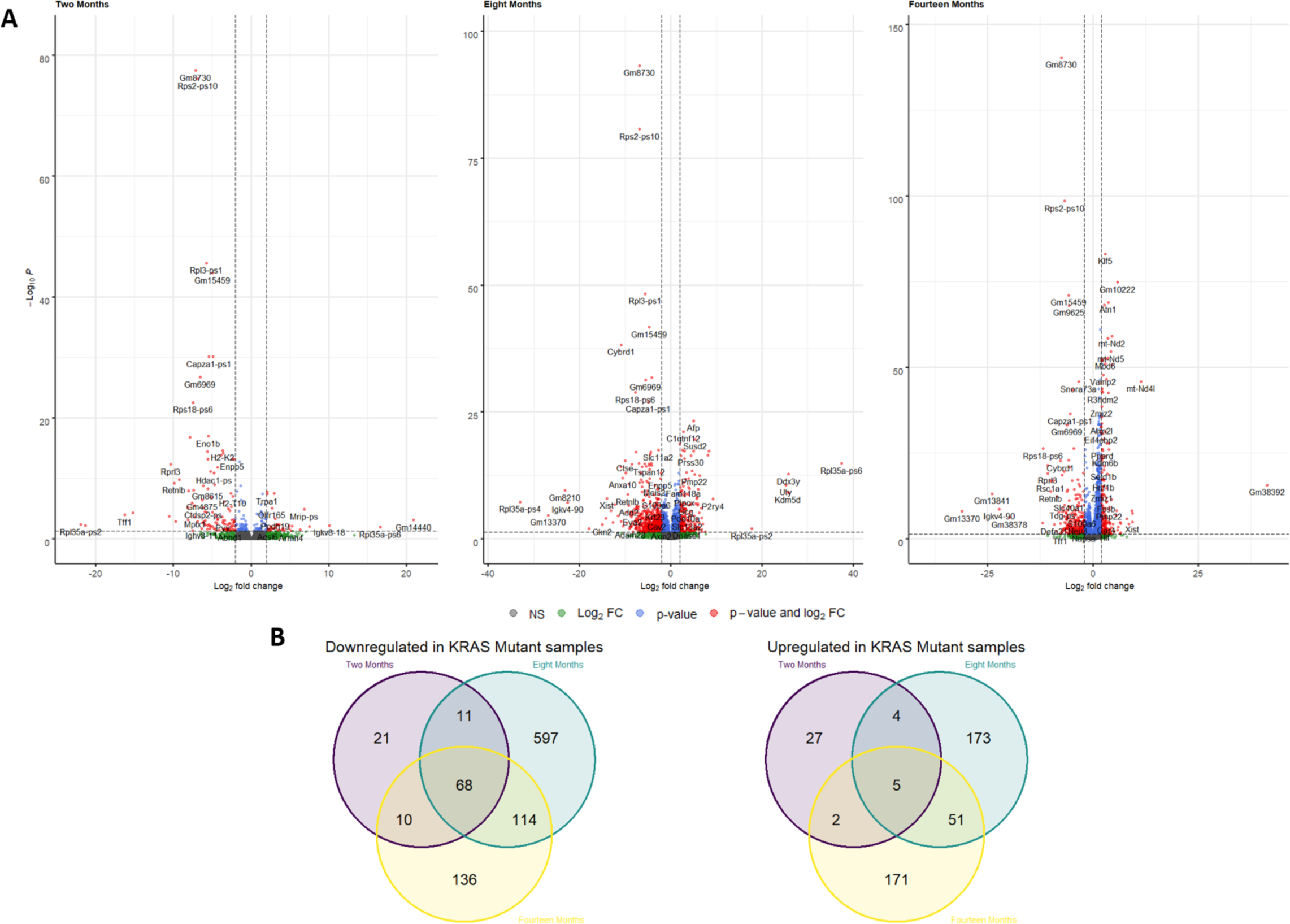
A) Differential gene expression analysis comparing the expression profiles of *Braf* and *Kras* mutant at two, eight, and fourteen months. B) Venn diagrams displaying the overlap between differentially expressed genes at two, eight and fourteen months.

**Table 1:** Differentially expressed genes in comparisons of the Kras and Braf mutant intestine at two, eight, and fourteen months. Differential gene expression analysis was performed using the DEseq2 R package. Genes with an absolute logFC > 2 and an FDR < 0.05 were deemed significant.

| Timepoint | Number of Genes |  | Sum |
| --- | --- | --- | --- |
|  | Overexpressed<sup>1</sup> | Underexpressed<sup>1</sup> |  |
| 2 Months | 38 | 110 | 148 |
| 8 Months | 233 | 790 | 1023 |
| 14 Months | 229 | 328 | 557 |

<sup>1</sup>Over (Under) expressed in *Kras* mutant samples relative to *Braf* mutant samples.

To explore the functional consequences of the disparate gene expression profiles of the *Kras* and *Braf* mutated intestine, we performed gene ontology enrichment analysis on genes differentially regulated at ≥ 2 time points. For genes that were downregulated in the *Kras* mutant intestine relative to *Braf* mutant animals, 203 were downregulated at ≥ 2 time points (Figure 4B – Venn Diagrams). This geneset was enriched for several immune related processes, including the immune response (Observed/Expected ratio: 2.84, FDR Corrected P=0.002), innate immune response (Observed/Expected ratio: 2.37, FDR Corrected P=0.003), and positive regulation of the immune response (Observed/Expected ratio: 3.84, FDR Corrected P=0.015). Other enriched processes include the metabolic process (Observed/Expected ratio: 1.51, FDR Corrected P = 0.041), and positive regulation of NF-kappa beta transcription factor activity (Observed/Expected ratio: 7.77, FDR Corrected P = 0.047). For genes upregulated in *Kras* mutant samples, 62 were upregulated at ≥ 2 time points, however enrichment analysis did not identify any biological processes associated with these genes.

### Immune cell deconvolution analysis reveals a *Braf-*mutation induced macrophage-inflamed response in the small intestine

Our earlier analysis revealed a strong immunological response in the *Braf* mutant intestine. We examined the immune contexture of the intestine of *Braf* and *Kras* mutant intestine by deconvoluting bulk RNAseq data using ImmuneCC (Fig 5A). Deconvolution analyses indicated a significant reduction in macrophage infiltrate in the microenvironment of the *Kras* mutant intestine when compared with *Braf* mutant mice (P=0.02, Fig 5B). The proportion of other immune cell types was not significantly different. We do note that some animals show a striking CD4+ T cell signature (Fig 5A), however this was not altered between *Braf* and *Kras* mutant animals.

**Figure 5:**
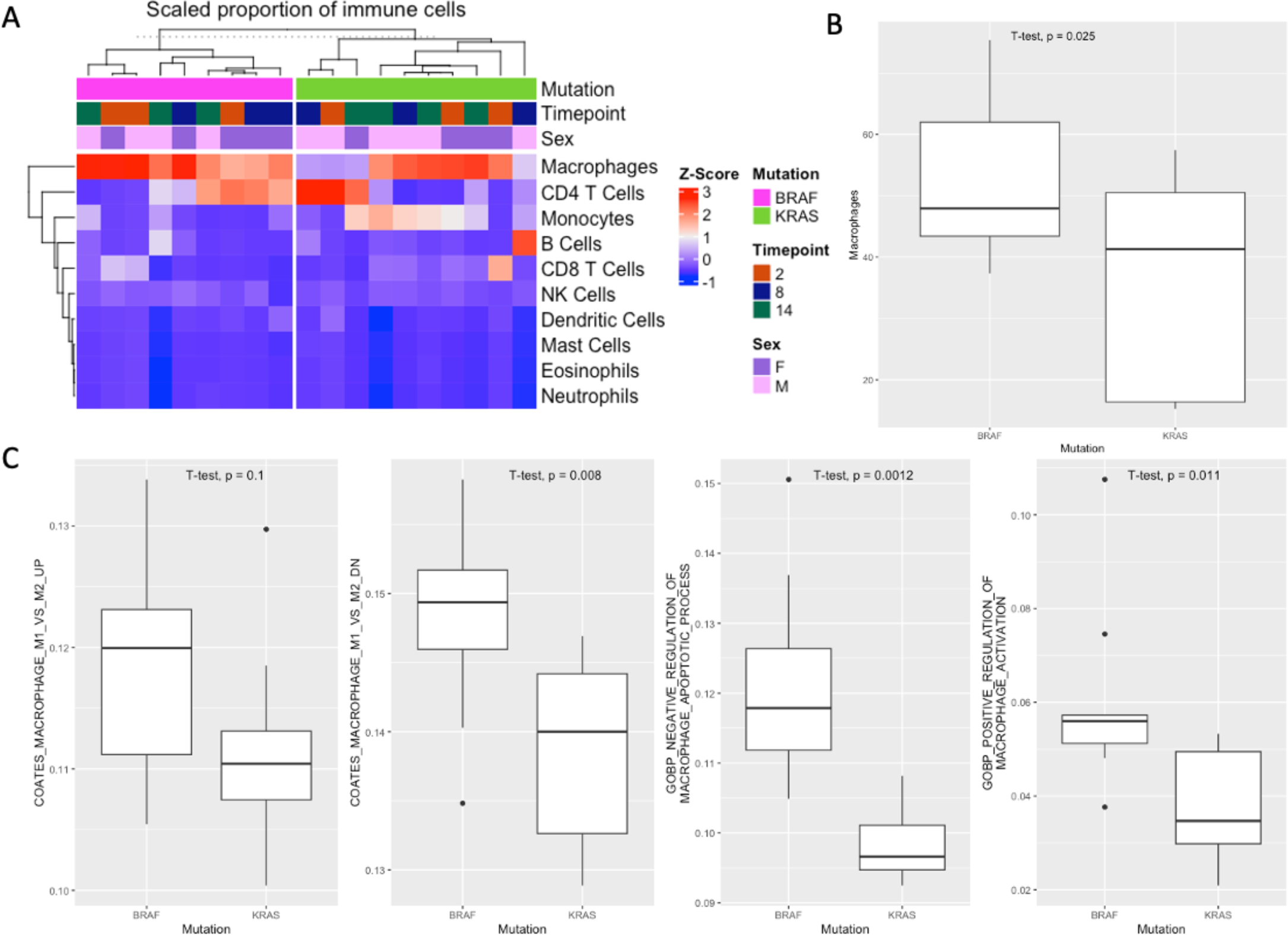
A) Relative abundance of immune populations in *Braf* and *Kras* mutant animals as measured via deconvolution of bulk RNA sequencing data. B) Relative abundance of macrophages in *Braf* and *Kras* mutant animals. C) Gene set enrichment analysis for gene sets involved in macrophage activation, and apoptosis.

Macrophages can be polarized into a M1 (pro-inflammatory) and M2 (anti-inflammatory state). To characterize whether *Braf* mutation could induce an M1 macrophage polarization phenotype we performed geneset enrichment analysis using previously published signatures from M1 and M2 macrophages. We observed significant enrichment for M1 inflammatory macrophage gene signatures in the *Braf* mutant model relative to *Kras* mutant mice (P=0.1 and P=0.008 for upregulated and downregulated genes that differentiate between M1 and M2 macrophages, respectively), in keeping with our observation that inflammatory processes and macrophages are enriched in the *Braf* mutant model (Fig 5C). We also observed a significant difference in enrichment scores for genesets associated with regulation of macrophage apoptosis (P=0.0012; Fig 5C) and macrophage activation (P=0.011; Fig 5C). These data indicate that M1 macrophages may be enriched in the hyperplastic intestinal mucosa of *Braf* mutant mice and may contribute to chronic inflammatory signals that drive lesion initiation.

### Intestinal Kras mutation is typified by an attenuation of the CpG island methylator phenotype

In human colorectal cancers both *KRAS* and *BRAF* mutation are linked to the CpG Island Methylator Phenotype, however we have previously reported differences in the methylation spectrum by mutation. We hypothesize that differences in the degree and extent of CIMP in these animals may explain the differential risk of transformation associated with each oncogene. To test this, we performed genome-scale DNA methylation analysis by reduced representation bisulphite sequencing (n=3 animals/genotype/timepoint; n=3 WT animals/timepoint).

First, we sought to explore the genome-wide implications of *Braf* and *Kras* mutation by performing principal components analysis (Figure 6A). PC1, explaining ∼10% of the variance in the data, was correlated with age in the wild-type animals. This represents “epigenetic drift”. In keeping with our earlier work, *Braf* mutant animals appear to accelerate endogenous aging, as evidenced by their accelerated divergence from wild-type animals across PC1 (Figure 6A). *Kras* mutant animals showed a similar acceleration of endogenous aging, however this was attenuated when compared with age-matched *Braf* mutant animals (Figure 6A).

**Figure 6:**
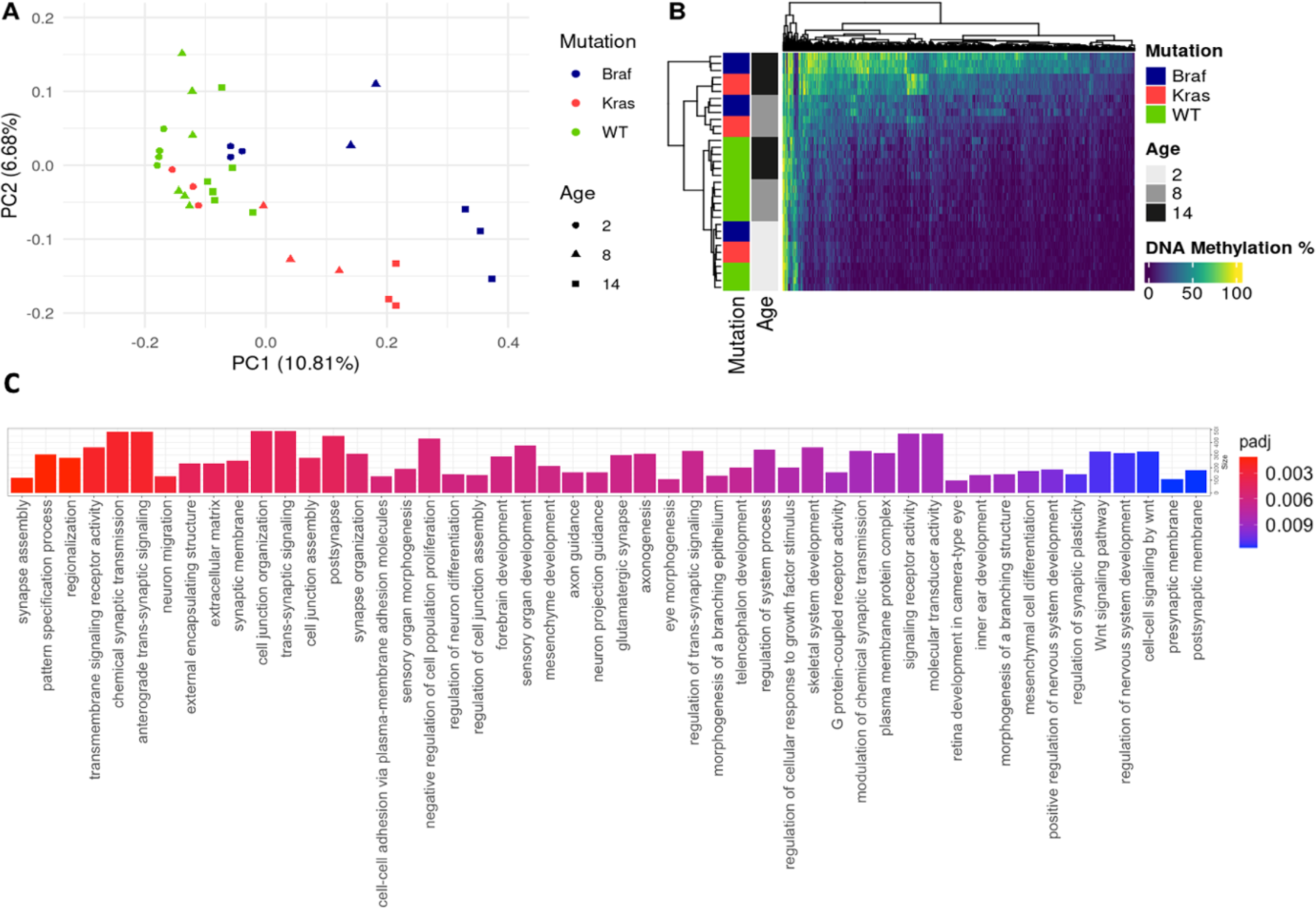
A) Principal components analysis of the entire methylome of *Braf, Kras* and WT animals. PC1 is strongly associated with age and exposure time to either *Braf* or *Kras* B) Heatmap depicting 1,306 CpG sites that show a significantly different temporal methylation trajectory between *Braf* and *Kras* mutant animals. C) Gene ontology analysis of differentially hypermethylated genes in *Braf* mutant animals compared with *Kras* mutant animals at 14 months.

To examine the interaction between each mutation and age, we performed ANCOVA analysis on CpGs associated with age. Of the 27,957 CpGs that were associated with age in *Braf* mutant animals (FDR<0.05), 55.1% were also associated with temporal shifts in DNA methylation in the intestine of *Kras* mutant animals. The rate of change in DNA methylation over time was significantly different in 1,306 of these sites (Figure 6B), with most sites accumulating DNA methylation at a slower rate in *Kras* mutant animals, providing further evidence for an attenuation of the CpG island methylator phenotype.

The discrepancy in the incidence of advanced serrated neoplasia between *Braf* and *Kras* mutant animals is most apparent at 14 months (Figure 1B). To understand the DNA methylation dynamics at this timepoint we performed differential DNA methylation analysis. We identified 2,231 differentially methylated CpG sites (FDR < 0.05, |Δ| methylation > 20), of which 1,803 and 428 were comparably hypermethylated and hypomethylated in *Braf* mutant mice, respectively. Gene ontology enrichment analysis of genes that are comparably less methylated in the *Kras* mutant mice compared with *Braf* mutants revealed 81 significantly enriched terms (Figure 6C). This included terms associated with differentiation, response to growth factors, WNT signalling, proliferation and cell-junction formation. Collectively, these data indicate that while *Kras* mutation induces widespread DNA hypermethylation, the severity and extent of methylation is lower when compared to that induced by *Braf* mutation.

## Discussion

Both *BRAF* and *KRAS* are important oncogenic drivers in multiple cancer types including colorectal cancer. Mutation of these genes is common in colorectal cancer and both drive overactivation of the MAPK signalling pathway, resulting in heightened proliferative activity, dysregulated differentiation, and evasion of apoptosis. They are mutually exclusive presumably because they act within the same pathway. Here, we have comprehensively assessed the propensity for mutation in *Braf* and *Kras* to induce spontaneous neoplastic transformation in murine models. The penetrance of neoplasia was substantially lower in *Kras* mutant mice, and invasive cancer formation was rare. We observe a striking association between neoplasia and gender in *Kras,* but not *Braf* mutant mice, with neoplasia more likely to occur in male animals. Our transcriptomic and DNA methylation analysis revealed differences in immune expression profiles and regulation of genes associated with key signalling axis and biological processes. We show a novel attenuation of the CpG island methylator phenotype and a distinct lack of WNT signaling dysregulation in *Kras* mutant mice. These findings shed light on differences in the natural course of polyps initiated by *Braf* and *Kras* mutation and highlight the restraints on the ability of mutant *Kras* to induce serrated pathway colorectal cancer.

Serrated neoplasia is hallmarked by overactivation of the MAPK signalling pathway (18), and widespread DNA methylation alterations (3,19,20). Hyperplastic polyps are common, small serrated polyps which are not considered to have significant malignant potential and can be initiated by either BRAF or KRAS mutations. The serrated lesions which have risk of malignant potential are sessile serrated lesions and traditional serrated adenomas. Sessile serrated lesions usually harbour *BRAF* V600E mutation, however the much rarer traditional serrated adenomas can activate MAPK via *KRAS* mutation in the context of additional genetic alterations which perturb BMP and WNT signalling (21–23). Interestingly, the 75% of colorectal cancers which arise from conventional adenomas initiated by *APC* mutations which perturb WNT signalling, often have *KRAS* mutations but very rarely *BRAF* mutations. In the context of conventional adenomas, *KRAS* mutations occur in advanced adenomas which are likely to transition to invasive cancers. It has never been fully understood why there is selective advantage for either *BRAF* and *KRAS* mutations in particular polyp types when both oncogenes activate the same pathway.

To examine the mechanics of *BRAF* and *KRAS* induced neoplasia, we induced the murine analogues of these mutations in the intestine of mice at weaning and monitored the temporal development of histological manifestations thereafter. Animals with *Braf* mutation spontaneously developed neoplasia with complete penetrance by 14 months, consistent with our prior work (17,24). By contrast, less than 50% of *Kras* mutant animals developed advanced neoplasia. This finding is consistent with previous literature indicating a limited capacity for *Kras* to induce intestinal neoplasia in mice (7). Feng and colleagues reported that *Kras* mutation induces overt proliferation and crypt hyperplasia, but not premalignant lesion formation or dysplasia. Our histological analysis confirms the former but disputes the latter. In our study, a subset of animals developed overtly dysplastic serrated lesions, however the *Kras* mutant animals in our study were aged to 18 months. These data indicate that the sojourn from mutation to lesion formation may proceed over a long period of time and that human *KRAS* initiated serrated neoplasia might follow a similarly protracted evolution.

We also examined the beta catenin/WNT signaling axis by immunohistochemistry. *Braf* mutant murine serrated neoplasia is predicated upon the spontaneous activation of WNT signaling via *Ctnnb1* mutation (8). Early *Braf* mutant lesions rarely mutate *Ctnnb1*, however by 14 months all animals displayed some form of *Ctnnb1* mutation. In humans, we observe mutation of *APC* or *RNF43* in dysplastic SSLs and serrated pathway cancers, with the former being more common in mismatch repair proficient cancers. *APC* is a member of the WNT pathway in keeping with our model system (6,25). Here, we report significant accumulation of beta catenin in lesions from *Braf,* but not *Kras* mutant mice. WNT activation is crucial for colorectal cancer progression and the lack of WNT activation in *Kras* mutant lesions contribute to their limited malignant potential.

The CpG Island Methylator Phenotype can occur in both *BRAF* and *KRAS* mutant colorectal cancers. CIMP has been linked to the silencing of key tumour suppressor genes, fueling cancer initiation and progression. We previously reported distinct subtypes of CIMP-High colorectal cancers, marked by differential mutational frequency of *BRAF* and *KRAS* (19). To evaluate the extent to which *Braf* and *Kras* mutation induce different DNA methylation patterns, we performed genome-scale methylation analysis on the intestine of mice. While both *Braf* and *Kras* mutant mice develop significant aberrant DNA methylation, the phenotype is less severe and somewhat attenuated in the *Kras* mutant intestine. This is consistent with the delayed development of neoplasia in these animals. *KRAS* mutation is common in human hyperplastic polyps and *BRAF* mutation is common in sessile serrated lesions. *KRAS* mutant hyperplastic polyps are rarely CIMP-high (26). By contrast, ∼70-80% of *BRAF* mutant sessile serrated lesions harbour the CpG Island Methylator Phenotype (27). Collectively, these data indicate that the attenuation and delayed development of CIMP may restrain *KRAS* mutant polyps and prevent their progression to cancer unless the KRAS mutation occurs in the context of pre-existing significant other mutations perturbing pathways such as WNT.

Despite inducing neoplasia with significantly lower penetrance, we observed elevated Ki67 expression in the crypt compartment of *Kras* mutant mice, indicating greater proliferative activity. This finding appears counterintuitive, but may be explained by the competition dynamics that underpin lesion formation in the gut. We and others have reported that WNT signalling is perturbed during aging (24,28,29), resulting in decreased stemness and regenerative capacity (30,31). We have also previously reported that WNT pathway members become hypermethylated during aging and that this process is hastened by *Braf* mutation (24). Flanagan and colleagues (32) recently reported on the role the competition dynamics in the intestine that allow for the formation of *Apc* mutant lesions. They show that NOTUM, a potent secreted WNT antagonist, inhibits WNT signaling in non-neoplastic crypts that neighbor lesions (32). This reduces their competitiveness and allows lesions to fix and expand. It is possible that the sustained elevation of proliferation in the *Kras* mutant crypts inhibits the formation of lesions by outcompeting lesions when they are initiated. In the *Braf* mutant intestine, which has comparably lower proliferation and stronger transcriptomic and epigenetic repression of WNT signaling and stemness, new lesions may have a competitive advantage and be able to form more readily. The precise role of competition dynamics in the serrated pathway warrant further investigation.

Our transcriptomic analyses revealed several genomic differences between the *Kras* and *Braf* mutated intestine. We note that the *Braf* mutant intestinal transcriptome is characterized by upregulation of immunological pathways, consistent with an inflammatory phenotype. To understand this we performed deconvolution analysis, revealing a significantly higher contribution of macrophages to the overall transcriptomic profile of these animals. Tissue resident macrophages can adopt pro- or anti-inflammatory phenotypes (33). Gene set enrichment analysis confirmed an upregulation of pro-inflammatory M1 macrophage gene expression programs, and concomitant decrease in apoptosis and increase in macrophage activation. Whether M1 macrophages promote or suppress neoplasia is context dependent. In established tumours, M1 macrophages are generally associated with tumour suppressive capabilities, owing to their secretion of pro-inflammatory cytokines which can enhanced tumour killing (34,35). By contrast, in inflammatory disorders they may promote tumour initiation by fostering a mutagenic microenvironment and promoting the generation of reactive oxygen species (36,37). It is possible that chronic inflammatory signaling, induced by increased abundance of M1 macrophages, may damage the epithelium and contribute to contribute to the increased propensity for neoplastic transformation in the intestine of *Braf* mutated animals.

Here we performed a comprehensive analysis of the impacts of both *Braf* and *Kras* mutation on the intestinal epithelium using transgenic murine models. We show that *Braf* mutation induces serrated neoplasia with high penetrance. This is accompanied by temporal dysregulation of the transcriptome and epigenome. We observed functional enrichment of immune related processes in the *Braf* but not the *Kras* mutated intestine. Immune deconvolution analysis provides evidence for a role of pro-inflammatory macrophages in the *Braf* mutated intestine. By contrast, *Kras* mutation does result in premalignant lesion formation, but this occurs over a significantly longer period time. Transcriptomic and DNA methylation alterations are attenuated in comparison to *Braf* mutant miceMurine serrated lesions in the *Kras* mutant intestine do not display hallmarks of WNT activation, and this may serve as a block to their progression. These data shed light on the differential role of *Braf* and *Kras* mutation in the gut and provide clarity around why *Braf* mutation induces serrated pathway cancers, while *Kras* mutant serrated cancers are rare.

